# Loss of integration of brain networks after sleep deprivation relates to the worsening of cognitive functions

**DOI:** 10.1101/2020.07.15.197590

**Authors:** Pesoli Matteo, Rucco Rosaria, Liparoti Marianna, Lardone Anna, D’Aurizio Giula, Minino Roberta, Troisi Lopez Emahnuel, Paccone Antonella, Granata Carmine, Sorrentino Giuseppe, Mandolesi Laura, Sorrentino Pierpaolo

**Affiliations:** Department of Motor Sciences and Wellness, University of Naples “Parthenope”, Italy; Institute of Applied Sciences and Intelligent Systems, CNR, Pozzuoli, Italy; Department of Social and Developmental Psychology, University of Rome “Sapienza, Italy; Department of Applied Clinical Sciences and Biotechnology, University of L’Aquila, Italy; Institute for Diagnosis and Cure Hermitage Capodimonte, Naples, Italy; Department of Humanities Studies, University Federico II, Naples, Italy; Institut de Neurosciences des Systèmes, Aix-Marseille Université, Marseille, France

**Keywords:** Magnetoencephalography, brain network topology, cognitive function, attention, sleep deprivation

## Abstract

The topology of brain networks changes according to environmental demands and can be described within the framework of graph theory. We hypothesized that 24-hours long sleep deprivation (SD) causes functional rearrangements of the brain topology so as to impair optimal communication, and that such rearrangements relate to the performance in specific cognitive tasks, namely the ones specifically requiring attention. Thirty-two young men underwent resting-state MEG recording and assessments of attention and switching abilities before and after SD. We found loss of integration of brain network and a worsening of attention but not of switching abilities. These results show that brain network changes due to SD affect switching abilities, worsened attention and induce large-scale rearrangements in the functional networks.

## Introduction

Historically, behavioral functions have been associated to localized brain structures (e.g. the Broca area for language production). More recently, converging evidence has showed that more complex, associative functions cannot be as easily associated to localized areas, but rather need distributed coordinated activity across multiple, diffuse areas (McGonigle et al., 2000, Ascoli et al., 2004). Hence, complex functions such as attention, memory and reasoning are modulated by a “multifocal neural system” formed by multiple interacting areas (McIntosh, 2000; Mesulam, 1990). As a consequence, capturing the complex patterns of coordinated activity is necessary in order to explain higher cognitive functions. To this regard, mathematical tools such as “Graph Theory” land themselves nicely to the study of the brain (Sporns, 2018). Under this framework the brain is conceptualized as a complex system of interacting units (Medaglia et al., 2015), represented as nodes (brain areas), linked by edges (representing structural, functional or causal interactions) (Bullmore & Sporns, 2009; Friston, 2011). This way, some topological features of the large-scale connections have been described consistently, such us “small-worldness” (Watts & Strogatz, 1998) and “scale-freeness” (Barabási & Albert, 1999). Such topological structure allows an optimal and efficient communication under spatial and energetic constraints. Consequently, pathological conditions that disrupt this optimally tuned configuration are deleterious, either as excessive disconnections as in Alzheimer disease (Heringa et al., 2014; Jacini et al., 2018) or in terms of hyper-connectedness, as it occurs in epilepsy and amyotrophic lateral sclerosis (Sorrentino et al., 2018). In fact, while, on the one hand, disconnectedness hinders efficient communication, hyper-connectedness, on the other hand, impairs brain flexibility (Sorrentino et al., 2020).

Sleep is an active physiological process during which the body recovers, and it is essential for the well-being and for optimal behavioral and cognitive performance. For example, both non-REM (NREM) and REM sleep promote learning and memory (Klinzing et al., 2019; Tononi & Cirelli, 2014). NREM sleep is relevant to many cognitive functions (Anderson & Horne, 2003; Lafortune et al., 2014; Walker, 2009; Wilckens et al., 2016, 2017). For example, modulations in amplitude of low-frequency slow wave activity (SWA) during NREM sleep is associated with improved executive functions (Wilckens et al., 2016), and memory consolidation (Staresina et al., 2015). Wilckens showed that higher delta EEG power (0.5–4.0 Hz), particularly within the NREM period, has been associated with better working memory, set-shifting, and reasoning in older adults (Anderson & Horne, 2003; Wilckens et al., 2016). Furthermore, Clark et al. (1998) reported that the relative power in the low frequencies in the frontal EEG channels during the NREM phase was directly related to the performance in neuropsychological tests assessing prefrontal functions.

Accordingly, insufficient or poor sleeping negatively affects mood and cognition, and the effects of sleep deprivation (SD) are evident at both the behavioral and physiological level. In fact, SD affects memory, executive functions and attention (Tomasi et al., 2009; Chee & Chuah, 2008; Lim & Dinges, 2012). In particular, SD selectively impairs top-down, but not bottom-up processing (Gevers et al., 2015), two mechanisms underlying stimuli perception and interpretation (Moore & Zirnsak, 2017). On the one hand, top-down processing is internally induced, and information is sought out voluntarily, while bottom up processing is driven by the salient features of the stimuli (Katsuki & Constantinidis, 2014).

EEG- and magnetic resonance-based evidence indicate that sleep loss affects primarily the frontal lobes. For instance, studies have shown that sleep pressure (i.e. the felt urge to sleep) relates to increased theta power density mostly in the frontal areas (Cajochen et al., 2001; Finelli et al., 2000).

The effects of SD on brain connectivity have been investigated, reporting reduced connectivity within the dorsal attention network, the visual network and especially the default mode network (DMN) (Gao et al., 2015; Kaufmann et al., 2016; Yeo et al., 2015). A prolonged wakefulness acts against the physiological decoupling between DMN and attentional networks (Fox et al., 2005), and this feature is associated with the worsening of cognitive performances. In particular, impairment of vigilance and attention (Lim & Dinges, 2008) are related to the alteration of the changes in the salience and attentional network (Ma et al., 2015).

While fMRI offers optimal spatial resolution, it is suboptimal in terms of temporal resolution. Furthermore, the estimation of functional connection in fMRI is done by correlating the amplitudes of the signals. On the contrary, EEG has optimal temporal resolution but poor spatial resolution, given that the electric fields are distorted by the layers surrounding the brain. To that regard, Magnetoencephalography (MEG) is a useful neurophysiologic technique as it allows an optimal trade-off between spatial and temporal resolutions (Baillet, 2017; Lopes da Silva, 2013).

Based on source-reconstructed MEG signals, we built a functional network where the links represent synchronization between any two areas. To estimate synchronization, we choose the phase linearity measurement (PLM), a purely phase-based metric designed to be robust to noise and insensitive to volume conduction (Baselice et al., 2019). In order to avoid potential biases induced by different edge densities or average degree (van Wijk et al., 2010), we used the minimum spanning tree (MST; Tewarie et al., 2015), obtaining comparable networks (Tewarie et al., 2015).

To the best of our knowledge, this is the first MEG-based study on SD. Therefore, we present an experimental design to evaluate the physiological and behavioral effects of 24-hour long SD. We hypothesized that SD modifies the organization of brain oscillatory activity, as captured by modified functional topology, to a less efficient large-scale communication. We further hypothesize that such topological rearrangements would relate to cognitive performance. To test these, thirty-two young adults underwent MEG recordings and assessments of attentional functions and switching abilities before and after 24h of SD. Observed topological changes were then correlated to cognitive performance.

## Material and Methods

### Participants

Thirty-two young adult males were recruited (mean age ± SD, 24,84 ± 2,85 years). All participants were right-handed, Italian speaker and none of them had a history of medical, neurological or psychiatric illness nor of medication or drug intake. Requirement of inclusion were normal sleep duration and no excessive daytime sleepiness. The quality and quantity of the participants usual sleep and daytime sleepiness were assessed by Pittsburgh Sleep Quality Index (PSQI; Buysse et al., 1989), the Epworth Sleepiness Scale (ESS; Johns, 1991) and the Karolinska Sleep Diary (KSD; Akerstedt et al., 1994). Subjects with scores less than 5 on the PSQI and less than 7 on the ESS were included in the study. From the third day before the beginning of the experimental procedure each participant was required to maintain a regular sleep-wake cycle and actigraphic recordings (wActiSleep-BT, ActiGraph-BT, Pensacola, Florida) were collected to control the subjects’ compliance. The intake of coffee, beverages containing stimulating active ingredients and intense physical activity were prohibited starting 24 hours before the experimental procedure, which was performed during the working week to avoid changes related to weekend activities. All participants gave their written informed consent. The study was approved by Ethical Committee of Psychological Research of the Department of Humanities of the University of Naples Federico II (n prot 11/2020) and was conducted in accordance with the Declaration of Helsinki.

### Procedure

The procedure was carried out by 4 participants at a time so that the night of SD was spent in group. Experimental protocol included two sessions which took place at 09.00 a.m. at day 1 (T0) and 24 hours later at day 2 (T1). In each session the participants underwent MEG recordings at rest and immediately after they performed the Letter Cancellation Task (LCT) and the Task Switching Task (TS). The order of TS and LCT was randomized. During experimental sessions, participants were seated on a comfortable chair in a soundproof room. After the first session the participants were free to return to their daily life activities. At 09.00 p.m. they returned to the laboratory to begin the sleep deprivation night under the supervision of the experimenters. To prevent them from falling asleep they were allowed to take short walks outside the laboratory during the night. In both sessions, we assessed the perceived subjective state of sleepiness through the administration of the Karolinska Sleepiness Scale (KSS; Akerstedt et al., 1994), and the cognitive load by means of the NASA Task Load Index (NASA; Hart, 2006).

### Cognitive assessment

#### Letter Cancellation Task (LCT)

The Letter Cancellation Task (M. Casagrande et al., 1999; Maria Casagrande et al., 1997) requires participants to search and sign sequentially (from left to right and from top to bottom), as fast and as accurately as possible, three target letters within a 36 × 50 matrix of capital letters (fonts: New York, “12”) printed on an A4 paper sheet. A maximum completion time of 5 minutes was allowed. Every target appeared 100 times in a random sequence; for each matrix, 300 hits were possible. To avoid learning effects, different parallel forms with different target letters were used. The following dependent variables were considered: number of hits (as measures of accuracy) and number of rows completed (as index of speed).

#### Task Switching (TS)

In task-switching, two different tasks were performed in rapid succession and according to a random sequence of task presentation, so that the task to be executed might change from one trial to the next (“switch” trial), or be repeated (“repetition” trial). Task switches are usually slower and less accurate than task repetitions, and this difference is often referred to as the “switch cost” (SC). This cost is thought to reflect the time needed for the executive control processes to reconfigure the cognitive system for the execution of a new task (Rogers & Monsell, 1995; Rubinstein et al., 2001) so that it can be considered an operational measure of the executive control (Couyoumdjian et al., 2010). This test has been usually used in neuroimaging study on SD (Nakashima et al., 2018).

All the participants were individually tested in a well-lit, sound-proof room. They were seated in front of a 15-inch computer monitor, at a distance of 50 cm, and at the beginning of each session, task instructions were both displayed on the screen and explained verbally by the experimenter, emphasizing the need for both accuracy (avoiding errors) and speed (minimize reaction times). In this study, the two tasks require deciding if a digit stimulus was odd or even (task A), or if it was greater or smaller than 5 (task B). In each trial of the two tasks, a cue (the “square” or “diamond” respectively) indicated the specific task (A or B) to perform on the subsequent target stimulus that appeared inside the cue. Experimental subjects used their left and right index fingers to provide your response: odd digits and digits smaller than 5 were mapped onto the left index finger response, whereas even digits and digits larger than 5 were mapped onto the right index finger response. The same two response keys on the computer keyboard (‘A’ for left and ‘L’ for right index finger) were used for both tasks. Stimuli presentation and response recordings were managed to employ custom software (Superlab, version 4.0.4 for Windows, Cedrus Corporation).

Each participant initially performed a training session (2 block of 80 trials) followed by an experimental session consisting of 320 trials, arranged in 4 blocks of 80 trials each. We considered the task as learned when, during the training session, at least 85% of the correct response were achieved. On each trial, a cue was presented for 1000 ms, and then it was followed by a target stimulus that remained on the monitor until the participant’s response. A schematic description of the present task-switching paradigm is reported in Figure 1.

**Fig 1.**
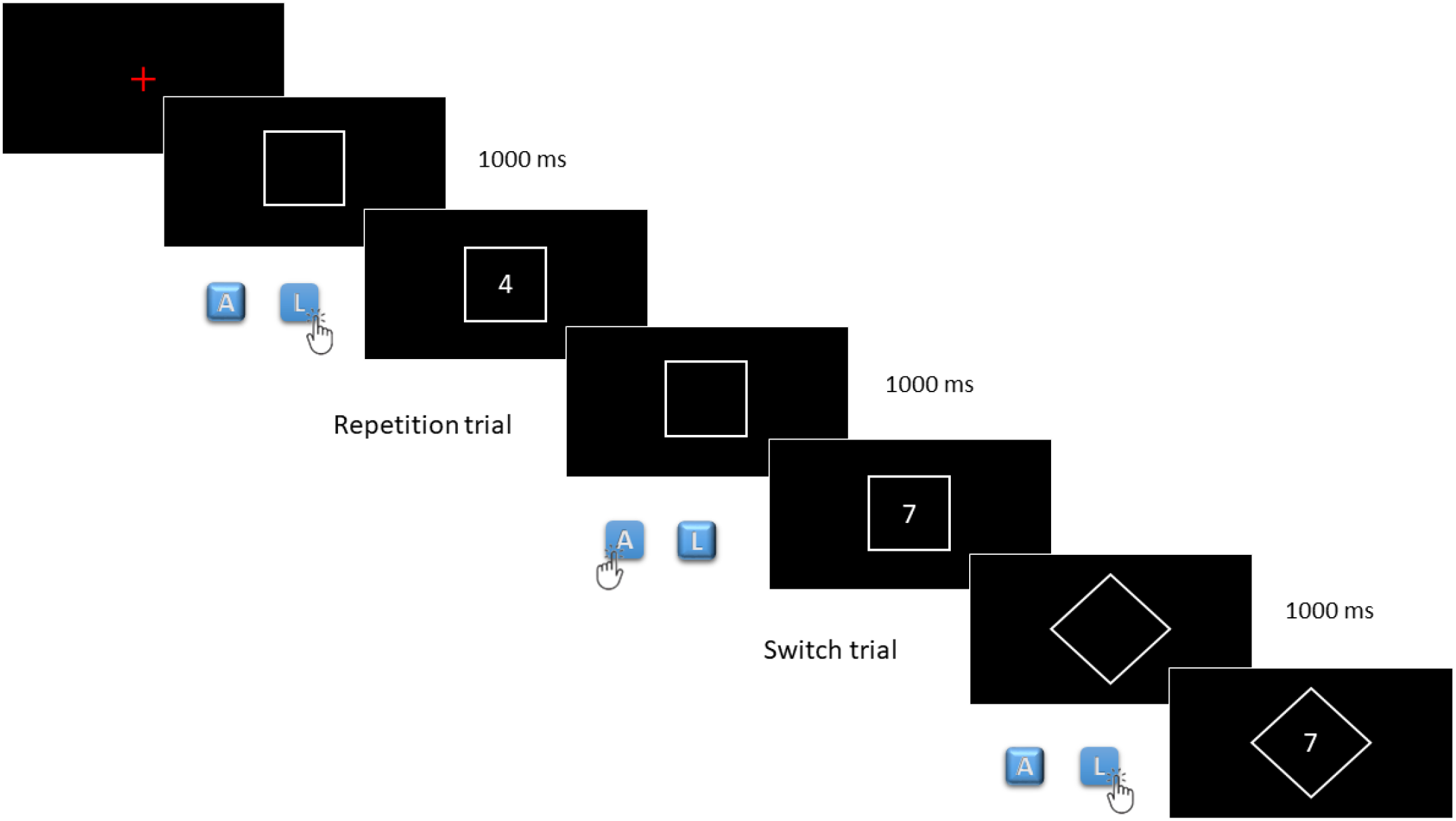
Schematic representation of Task Switching. The two tasks require deciding if a digit stimulus was odd or even (task A), or if it was greater or smaller than 5 (task B). In each trial, a cue (“square” or “diamond”) appeared first, indicating the task (A or B) to be performed on the subsequent target stimulus appearing inside the cue itself. The “L” key (right index finger) of the keyboard key was used provide the response “even” or “larger than 5”, while the “A” key (left index finger) was used to respond “odd” or “less than 5”. Each participant initially performed a training session (1 block of 80 trials) followed by an experimental session consisting of 320 trials, arranged in 4 blocks of 80 trials each. On each trial, the cue was presented for 1000 ms, and then it was followed by the target stimulus.

### MEG acquisition

The MEG system, developed by the National Research Council, Pozzuoli, Naples, at Institute of Applied Sciences and Intelligent Systems “E. Caianiello” (Granata et al., 2018; Rucco et al., 2019), is equipped with 154 magnetometers and 9 reference sensors located on a helmet. The system is placed inside a room shielded from external magnetic fields (AtB Biomag UG - Ulm – Germany). Before signals recording, four position coils were placed on the participant’s head and digitized by using Fastrak (Polhemus®). The coils were activated and localized at the beginning of each segment of registration. The magnetic fields were recorded for 7 minutes, divided into two time intervals of 3’30”, while the participants were sitting comfortably in an armchair in the cabin with their eyes closed without sleep. Before the acquisition, for each participant, a standardized instruction to stay with eyes closed was communicated through intercom. Electrocardiogram (ECG) and electro-oculogram (EOG) signals were also recorded during the acquisition. The data were sampled at sf=1024 Hz, and a pass-band filter between 0.5 and 48 Hz was applied offline, during data pre-processing after the recording phase.

After the recording phase, the time-series were cleaned through an automated process (Sorriso et al., 2019). The FieldTrip toolbox (Oostenveld et al., 2011), which is based on Mathworks^®^ MATLAB^™^ environment, was used to implement principal component analysis (PCA), to reduce the environmental noise, and independent component analysis (ICA), to remove the physiological artefacts such as cardiac activity or eyes blinking (if present). For each participant, the cleaned trials longer than 4s were selected. On the latter, a beamforming procedure was performed using the Fieldtrip toolbox. Based on a magnetic resonance imaging (MRI), the volume conduction model proposed by Nolte was considered and the Linearity Constrained Minimum Variance (LCMV) beamformer was implemented to reconstruct time series related to the centroids of 116 regions-of-interest (ROIs), derived from the Automated Anatomical Labeling (AAL) atlas. Subsequently the Phase Linearity Measurement (PLM) was performed to provide an adirectional estimate of the connectivity that is purely based upon the phases of the signals, and insensitive to volume conduction. The PLM is defined by the following equation:

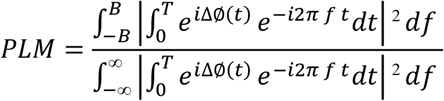

where the Δø(*t*) represent the phase difference between two signals, the 2B is the frequency band range, *f* is the frequency and *T* is the observation time interval. The PLM ranges from 0 to 1, where 1 entails perfect synchronization, and 0 conveys the absence of phase relationship.

By computing PLM for each couple of brain regions, we obtained a 90×90 weighted adjacency matrix for each epoch and for each subject, in all of the frequency bands: delta (0.5–4 Hz), theta (4.0–8.0 Hz), alpha (8.0–13.0 Hz), beta (13.0–30.0 Hz) and gamma (30.0–48.0 Hz). We excluded from the analysis the cerebellar regions, given thei low reliability, leaving 90 regions encompassing the cerebral cortex and the basal ganglia. The weighted adjacency matrix was used to reconstruct a brain network, where the 90 areas of the AAL atlas are represented as nodes, and the PLM values form the weighted edges. For each trial longer than 4s and for each frequency band, through Kruskal’s algorithm, the minimum spanning tree (MST) was calculated. The MST is a loop less graph with N nodes and M= N-1 links. The MST was computed to obtain topologic measures that are unaffected by the degree distribution, matrix density, or arbitrary thresholds.

Finally, both global and nodal parameters were calculated. The global parameters included: the *diameter* (D), defined as the longest shortest path of an MST, representing a measure of the ease of communication across a network; the *leaf fraction* (L), defined as the fraction of nodes with degree equal to 1 (“leaf”), providing an indication of the integration of the network; the *degree divergence* (K), a measure of the broadness of the degree distribution, related to resilience against targeted attacks; and the *tree hierarchy* (Th), defined as the number of leaves over the maximal betweenness centrality, meant to capture the optimal trade-off between network integration and resiliency to hub failure. The nodal parameters included: the *degree* (k), defined as the number of connection incident on a given node; and the *betweenness centrally* (BC) described as the number of shortest paths passing through a given node over the total of the shortest paths of the network. Before moving to the statistical analysis, all the metrics were averaged across epochs in order to obtain one value for subject. A pipeline of the processing MEG data is illustrated in Figure 2.

**Fig 2.**
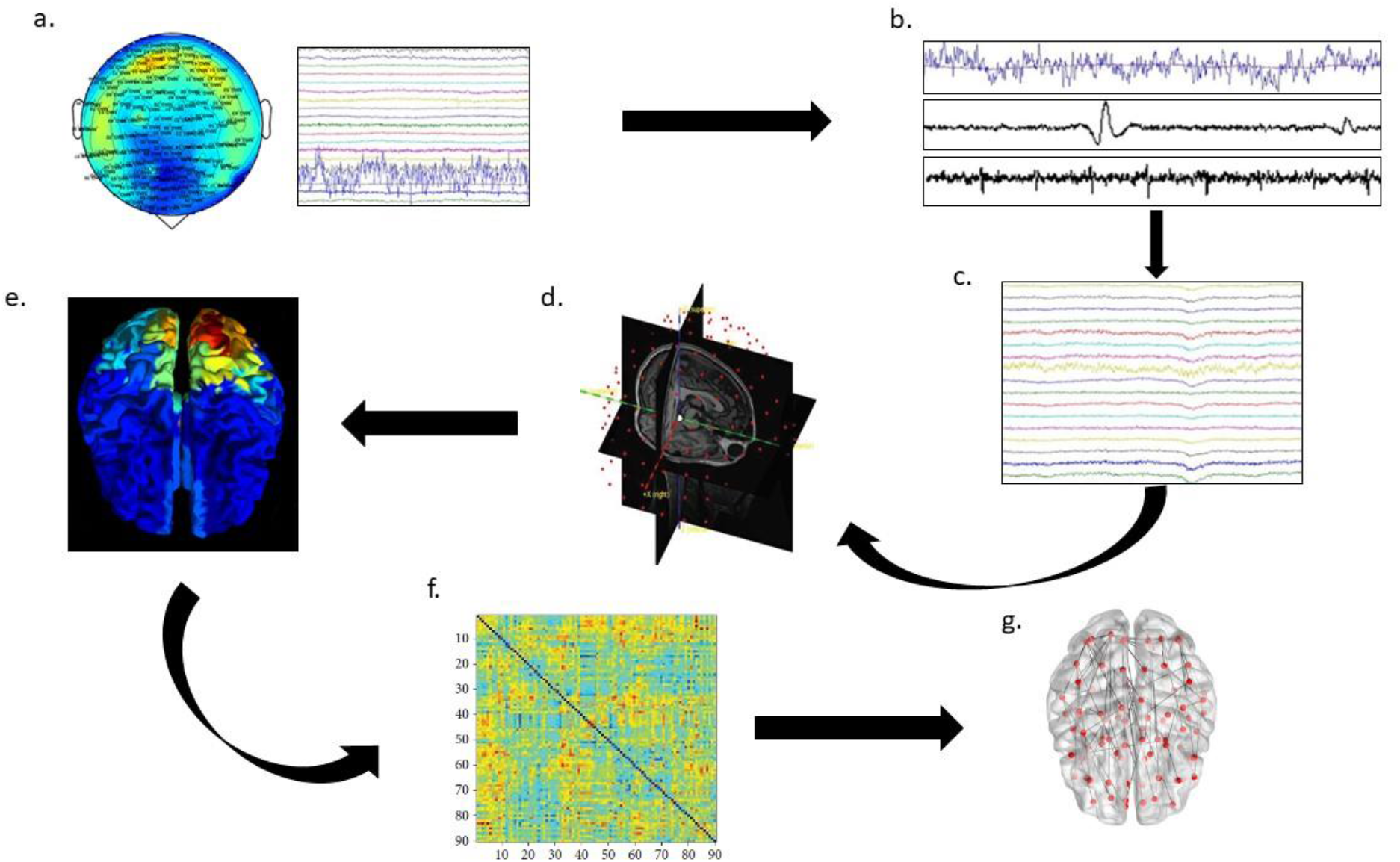
Data analysis pipeline: a) MEG signal recorded by 154 sensors; b) noisy channels, blinking and cardiac artifact are removed to obtain the cleaned signals; c) cleaned signals; d) MRI and MEG sensors are coregistered; e) the source activity has been estimated using a beamforming algorithm: the activity on the sensor level is projected onto brain regions (based on the AAL atlas); f) connectivity matrix: raws and colums are brain areas, entries are PLM values; g) brain network representation based on the MST.

**Fig 3.**
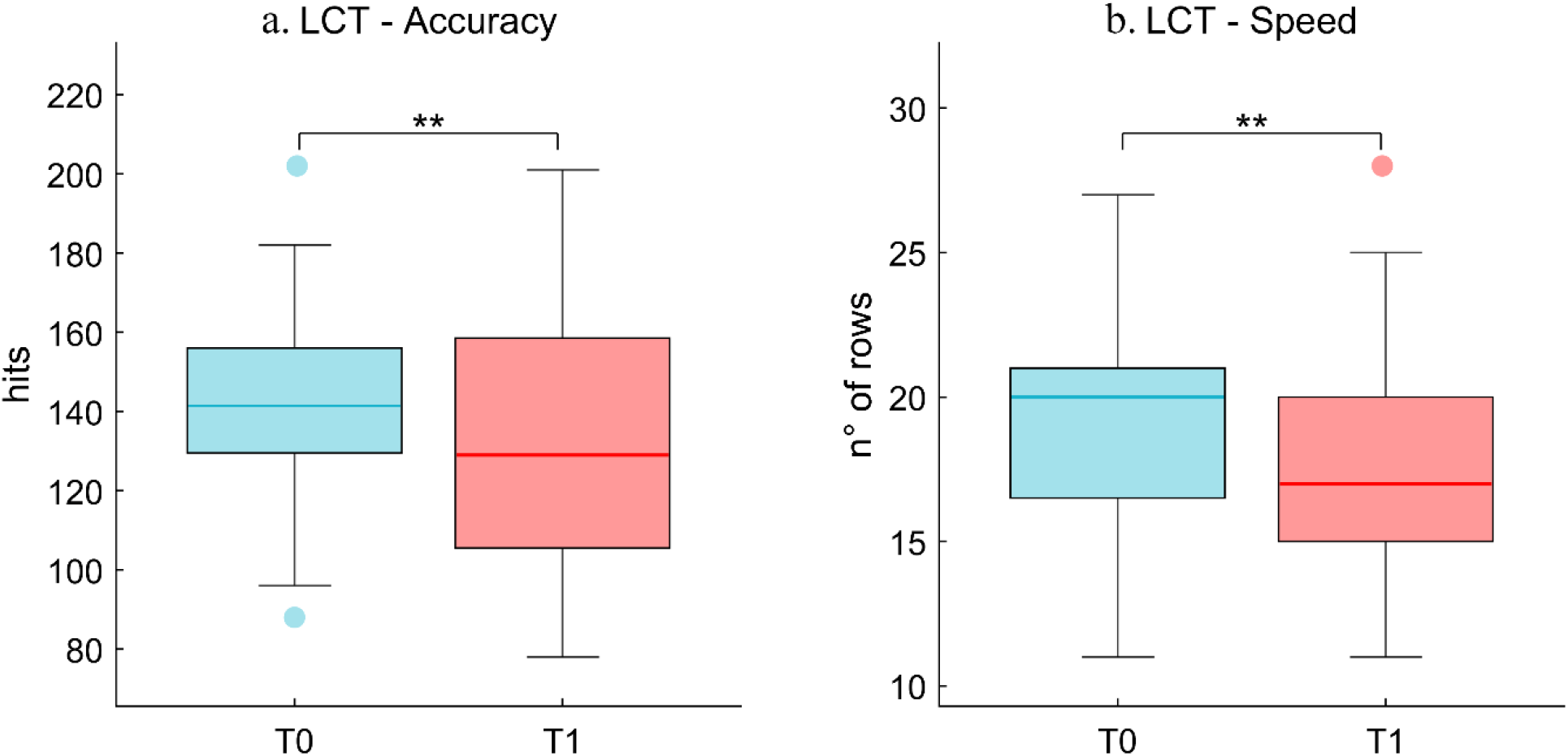
Letter Cancellation Test (LCT). Performance in LCT show the worsening of attentional funciton: accuracy (Fig.3a) and speed (Fig.3b) were reduced in T1 in respect to T0. ** = p<.01

**Fig 4.**
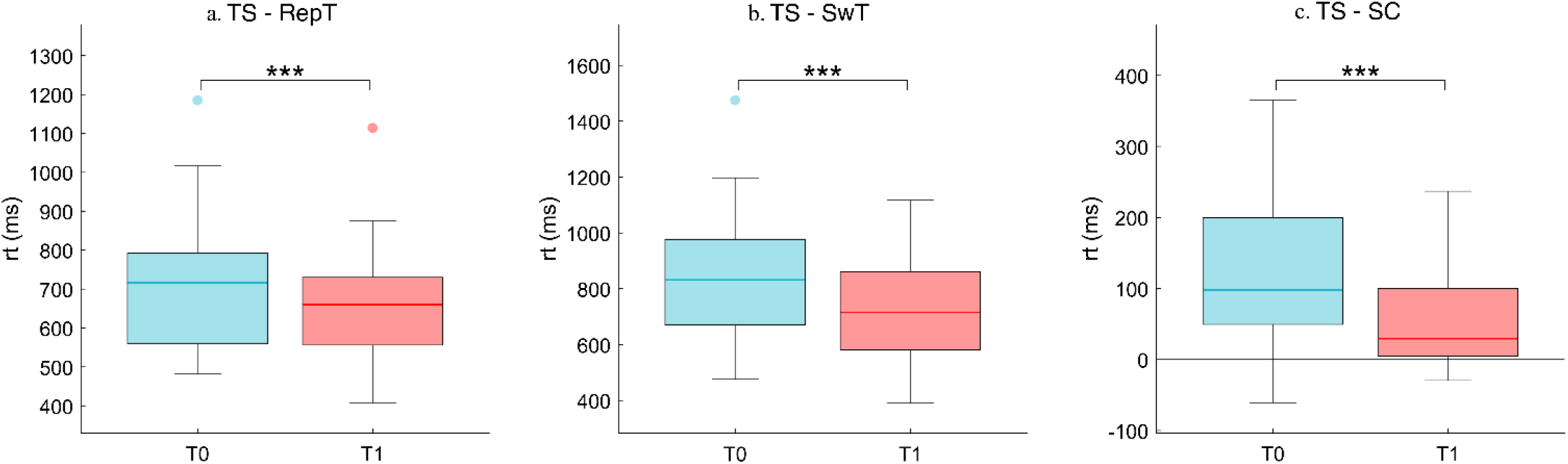
Task Switching. Task switching (TS) show a reduction in time reaction (rt) both in repetition trial (RepT; Fig.4a) and Switch trial (SwT; Fig.4b). Even the Switch cost (SC; Fig.4c) was reduced after 24h of sleep deprivation. *** = p<.001

**Fig 5.**
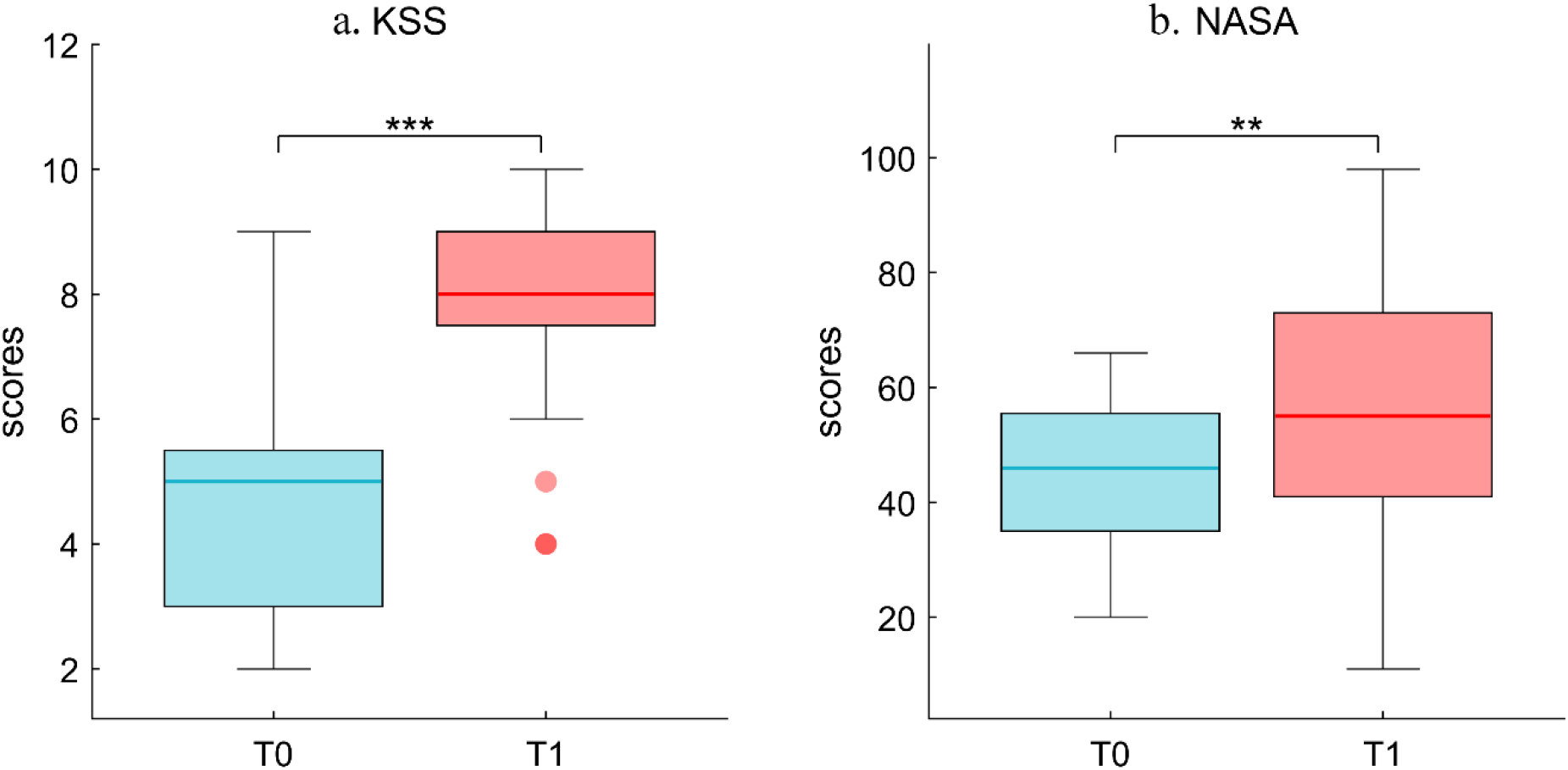
Subjective evaluation. Participants reported a significant increase in the subjective level of sleepiness (KSS; fig.3a) and cognitive load perceived (NASA; fig.3b) in T1 compared to T0. ** = p<.01, *** = p<.001

**Fig 6.**
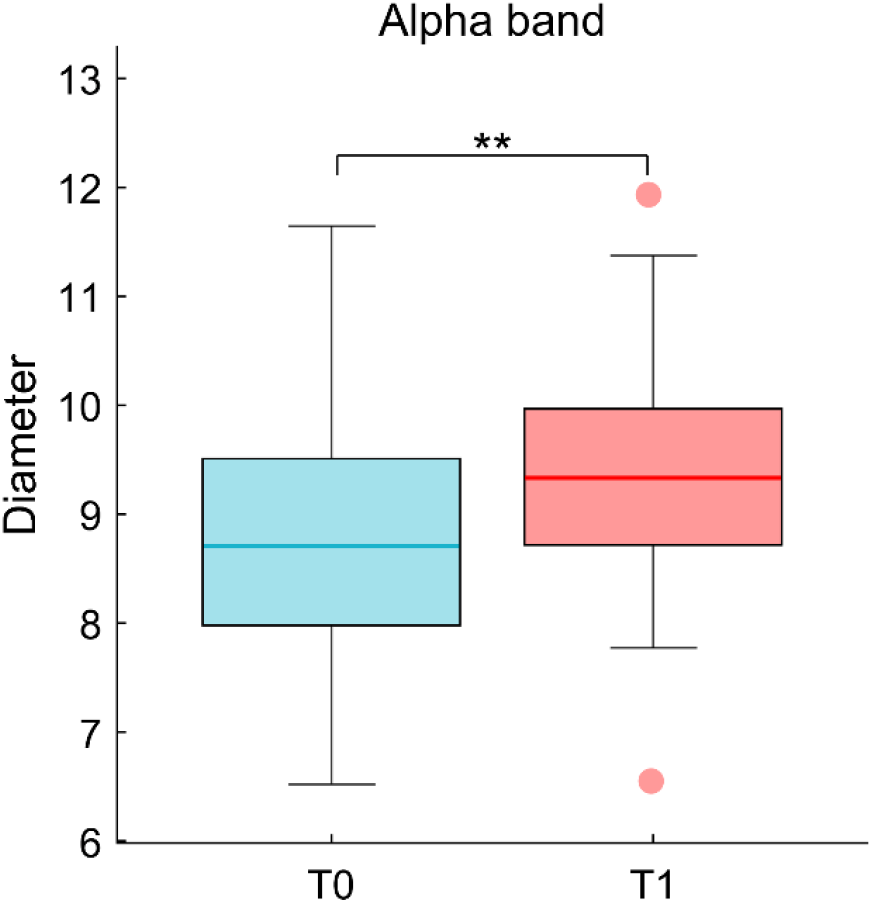
Topological analysis; Diameter. Statistical comparison between T0 and T1 show a significant increase in diameter in alpha bands after 24h of sleep deprivation. ** = p_fdr_<.01

**Fig 7.**
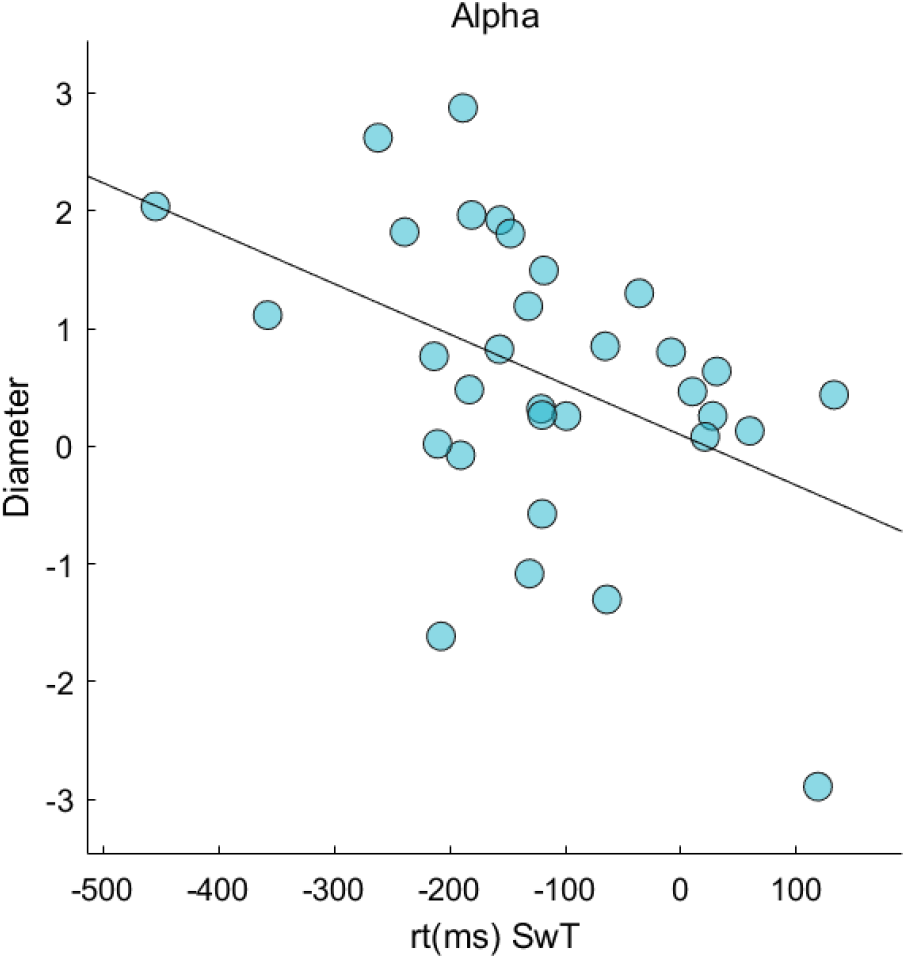
Correlation analysis. Pearson’s correlation between diameter in alpha band and Switch trial time reaction.

### MRI acquisition

MRI images of thirty-four young adult males were acquired on a 1.5-T Signa Explorer scanner equipped with an 8-channel parallel head coil (General Electric Healthcare, Milwaukee, WI, USA). MR scans were acquired after the end of the SD protocol. In particular, three-dimensional T1-weighted images (gradient-echo sequence Inversion Recovery prepared Fast Spoiled Gradient Recalled-echo, time repetition = 8.216 ms, TI = 450 ms, TE = 3.08 ms, flip angle = 12, voxel size = 1×1×1.2 mm1; matrix =256×256) were acquired.

### Statistical analysis

In order to assess the potential variations of the participants’ executive performance during the two experimental sessions (T0 vs T1), and to assess how both sleep deprivation and the consequent increase in sleepiness can affect their performance, we run two different statistical analysis, respectively two-factor repeated-measures ANOVA, with Time and Trial as within factor, and paired *t*-test.

In particular, with regards to the TS, median reaction times (in ms; median RT) to both repetition and switch trials, and angular transformations of the proportion of errors resulting from the two experimental sessions, were submitted to two-factor repeated-measures ANOVA. SC as well as all dependent variables obtained from LCT (number of hits and number of rows completed), were analyzed though paired *t*-test. SCs were computed as the difference between median switch RT and median repetition RT. Proportions of errors (EP) were computed by including both incorrect and missing responses. Before statistical analysis, this variable was submitted to an angular transformation, y=arcsen [sqr(p)], where sqr(p) is the square root of the proportion. All statistical analyses were performed using IBM SPSS Statistics for Macintosh, version 25.0 (IBM Corp., Armonk, NY, USA).

With regard to the topological data, statistical analysis was performed in Matlab (Mathworks®, version R2018b). Non parametric Wilcoxon test was performed to compare T0 and T1 in all frequency bands; all *p*-values were corrected for multiple comparison using the false discovery rate (FDR; (Benjamini et al., 1995)). Subsequently, the Pearson’s correlation index was used to find possible correlations between topological data and behavioral performances. We calculated the difference of the values of all the variables between T1 and T0. Therefore, the correlation analysis was carried out between the differences in topological parameters and the differences in scores on cognitive tests (LCT, TS) and subjective evaluation (KSS, NASA). The alpha level was fixed at 0.05.

## Results

### Cognitive assessment

#### Letter Cancellation Task (LCT)

In order to evaluate the attentional function, the LCT was administered. The participants were tasked to identify and mark 3 target letters within an array of distracting letters. Accuracy and speed in performing the test were evaluated. With regards the accuracy, a better performance (t_31_ = 2.8; p = 0.009) at T0 (mean ± se = 143.03 ± 4.32) than at T1 (mean ± se = 132.34 ± 6.02) was demonstrated (Fig.3a). With regard the speed, expressed as the number of inspected rows within the task time limit, a statistically significant difference was apparent between the two experimental sessions (t_31_ = 3.2; p = 0.003), as the subjects were faster in performing the test during T0 (mean ± se = 19.12 ± 0.63) with respect to T1 (mean ± se =17.75 ± 0.7) (Fig.3b).

#### Task Switching (TS)

In order to evaluate the switching abilities, the participants performed the TS task. This task requires to select the appropriate response based on the type of stimulus presented. Reaction times (RT), errors proportion (EP) and switch cost (SC) (as measure of executive control) were assessed.

A statically significant effect of time on RT (F_1,31_ = 15.82; p < 0.0001; ηp2 = 0.32) was observed, indicating, regardless of the trial, higher RT at T0 (mean ± se = 775.61 ± 36.22) than at T1 (mean ± se = 690.67 ± 28.32). There was also a statically significant effect of the trial on RT (F_1,31_ = 39.4; p < 0.0001; ηp2 = 0.54), revealing a faster RT in repetition trials (mean ± se = 689.2 ± 28.34) than in switch trials (mean ± se = 777.09 ± 36.09). Finally, a statically significant interaction effect was found between time and trial (F_1,31_ = 12.74; p = 0.001; ηp2 = 0.28; Repetition trials: T0 (mean ± se) = 715.32 ± 30.35; T1 (mean ± se) = 663.07 ± 25.83 (Fig.4a); Switch trials: T0 (mean ± se) = 835.9 ± 38.87; T1 (mean ± se) = 718.28 ± 30.22 (Fig.4b). With regards to EP, a statically significant effect of trial on EP (F_1,31_ = 8.79; p = 0.006; ηp2 = 0.21) was showed, indicating, regardless of the time, higher accuracy during the “repetition trials” (mean ± se =1.49 ± 0.01) with respect the “switch trials” (mean ± se =1.44 ± 0.02). No other significant effects were observed. Finally, significantly higher SC (t_31_ = 3,57; p = 0.001) at T0 (mean ± se = 120.57 ± 18.87) than at T1 (mean ± se = 55.2 ± 11.73) was observed (Fig.4c).

In order to evaluate a possible learning effect, before the first experimental session a training session was performed and a comparison between the second block of the training and the first block of the test was performed. Results showed a reduction of RT in switch trials (t_31_ = 3.6; p < 0.001; Training (mean ± se) = 1062.3 ± 60.2; Test (mean ± se) = 860.9 ± 58.7) and switch cost (t_31_ = 2.5; p < 0.05; Training (mean ± se) = 261.2 ± 36.6; Test (mean ± se) = 133.8 ± 37.8). However, no significant difference in EP (indicating the achievement of maximum performance during the trial session, before the test) was evident.

### Subjective evaluations

KSS and NASA have been used to assess the level of subjective sleepiness and perceived cognitive load. KSS results showed an increased level of sleepiness after 24 hours of sleep deprivation (Fig.5a; mean ± se; T0: 4.65 ± 0.32, T1:7.87 ± 0.25; *p* < 0.001), and the NASA index was increased in T1 as compared to T0 (Fig.5b; mean ± se; T0: 43.75 ± 2.24, T1:54.03 ± 3.72; *p* < 0.01).

### Topological analysis

The analysis for the global parameters showed increased Diameter in the Alpha band (Fig.6; median ± IQR; T0: 8,7 ± 1,48; T1: 9,33, 1,17; *p*_*fdr*_ < 0.01). No other global (leaf fraction, degree divergence, eccentricity and three hierarchy) nor nodal (degree, betweenness centrality) parameters were different between the two conditions.

### Brain topology, performance and sleepiness

In order to evaluate a possible relationship between network rearrangements and cognitive functioning, we carried out a correlation analysis between topological parameters, test performance and subjective evaluations. Specifically, Pearson’s correlation analysis showed a reverse correlation between the diameter in the alpha band and a Switch trial reaction time (rt SwT) (Fig.7; *p*_*fdr*_= 0.03, r = −0,447). The LCT performance, as well as the KSS and NASA scores, were not correlated with topological data.

## Discussion

SD is a typically harmful condition for the brain. With regard to moderate sleep deprivation, the main cognitive effects concern attentional functions and higher order cognition (Killgore, 2010). Moreover, SD is known to cause changes in neurophysiological activity (Krause et al., 2017).

In this paper, we set out to test the hypothesis that sleep deprivation causes functional rearrangements in the brain so as to impair optimal communication, and that such rearrangements relate to changes in specific cognitive tasks, especially the ones requiring sustained attention.

As we expected, after 24 hours of SD the participants reported increased subjective sleepiness, as seen from the scores of the KSS. SD negatively influenced attentional functions as evident from the worsening in the speed and accuracy of the LCT. Our data are in line with Casagrande et al., (1997) showing that the effect of SD on cognitive performance are prominent in tasks requiring high attentional load (both in short and long tasks). In our study, the impairment of attentional functions was demonstrated by the worsening in the three letter searching task. Accordingly, several authors (Lanteaume et al., 2016; Lim et al., 2010; Lim & Dinges, 2008) reported that attention is among the firsts functions to be affected by SD. Given the nature of the task, the attention deficit was possibly due to the weakening of the top-down processing, an internally induced process in which information is sought on voluntarily chosen factors (Katsuki & Constantinidis, 2014).

A counterintuitive scenario was observed in the TS where performance appeared slower at T0 than at T1, as shown by both higher RTs and SCs. Coherently, also SCs appeared to be reduced as a consequence of SD. This unexpected effect of a single night of sleep loss cannot be ascribed to data instability. In fact, as an indirect measure of data goodness, statistical comparison of reaction times and error proportions at task switching highlighted faster RTs and reduced EP in repetition than switch trials, as expected on the light of previous literature (Couyoumdjian et al., 2010), and this trend was confirmed in both pre- and post-sleep deprivation sessions. The unexpected improvement of the reaction times could lead erroneously to the conclusion that the switching ability improves following the SD. However, the lack of interaction effect on the errors committed between T0 and T1 shows that the performance was better only as regards the speed of response but not for the accuracy. In fact, the number of errors (a fundamental parameter to verify the learning of a task) remained unchanged between the two sessions, therefore it cannot be considered that the performance had improved in absolute terms. The hypothesis that these results may be due to a learning effect between the two sessions does not appear to be the most likely, as all participants reached the maximum level of performance during the training phase. In fact, a comparison between the second block of the training phase and the first block of the experimental session at T0 revealed the absence of differences in the number of errors, confirming that the task had been fully learned in the training phase. Thus, if on the one hand the test was found to be reliable, on the other end one cannot ignore the context in which the entire experimental procedure was carried out. An important role could have been played by the social component of the experimental setting as the SD was carried out in groups of 4 participants at a time (in addition to the presence of the experimenters). As known, most of the studies use forced wakefulness protocols to evaluate the effects of SD on cognitive function. However, in everyday life, people can voluntarily deprive themselves of sleep for various reasons such as work shifts and recreational activities. Multiple brain systems are affected differently based on the sleep deprivation model used. Deurveilher et al. (2013) showed that a condition of “voluntary” deprivation characterized by social interactions and exploration of the environment caused a greater activation of the cholinergic pathways in the basal forebrain compared to a “forced” deprivation characterized by simple sensory stimulation. In this way, the increased activation supported the motivation to stay awake by attenuating the signs of SD. Similarly, the social interaction condition experienced in our experimental setting may have contributed to the result obtained in task switching.

The topological analysis showed that after SD the brain network undergoes rearrangements at the global level. Global metrics allow to characterize the widespread remodeling of large-scale brain activity. We found increased diameter after SD in the alpha band. The diameter is defined as the longest shortest path of a graph (Tewarie et al., 2015). A network characterized by a small diameter presents a functional configuration with a few central nodes mediating long-range communication (Bullmore & Sporns, 2009; Van Den Heuvel et al., 2009; Van Den Heuvel & Sporns, 2011). This arrangement is advantageous as it allows efficient communication throughout the entire network, at the cost of increased risk of node overload (Stam, 2014). Conversely, a higher diameter implies a less compact topology with, on average, longer paths to go between any two nodes. This implies that the relative importance of nodes is more homogeneous (Rubinov & Sporns, 2010). Given the redistribution of the workload across the brain network, the node overload risk might be reduced. One possible interpretation of the increase in diameter that we observed would be that this topological rearrangement is in place to cope with cognitive demands after the SD, redistributing the computational load across the network. In fact, the inverse correlation between the diameter and the reaction times in switch trials of TS showed that topological rearrangements are related to cognitive performance. In particular, the sleep-deprived subjects have a less integrated topology and shorter reaction time. Hence, topological change allows to perform the task faster, specifically in switching trials (the most cognitively demanding trial). However, caution should be used when interpreting the data, given that the brain measures are made at rest and not during the execution of the task.

It is well known that a correct balance between integration and segregation of activity is relevant (Sporns et al., 2000). After 24 hours of prolonged wakefulness, the brain networks appeared to have modified its overall integration. The correct balance between integration and segregation seems to be mudulated by neurotransmitter-specific pathways. More specifically, cholinergic neurons have a strong influence in promoting segregation. Cholinergic tone is known to be modulated by wakefulness via the inhibition induced by adenosine (AD) on acetyilcholine (ACh) (Boonstra et al., 2007; Shine, 2019). In fact, during SD, the increase in AD causes a modulation of the cholinergic system that, from the basal forebrain, project widely towards subcortical and cortical structures including the visual, somatosensory and prefrontal cortex, controlling selective visual attention (Moore & Zirnsak, 2017; Noudoost & Moore, 2011). In addition to ACh, noradrenaline (NA) influences the network topology. Noradrenergic projections from the locus coeruleus are widespread to the whole brain and supports network integration (Shine, 2019). Furthermore, high levels of NA have been associated with the ability to shift between tasks(Aston-Jones & Cohen, 2005) and other higher order functions such as cognitive control. However, it is surprising that after 24h of SD the participants performed the TS faster. Our interpretation of the evidence is that this type of task is more engaging: in this case, the processing of information is externally driven through a bottom up mechanism. In fact, selective damage to top down versus bottom up mechanisms after sleep deprivation has been demonstrated by Gevers et al. (2015). Thus, the brain network was able to support optimal performance by compensating through greater cognitive effort, as suggested by the increased cognitive load detected through the NASA test. The increase in cognitive load compared to unchanged performance is in line with Tomasko et al., (2012) that showed that SD increases the cognitive load of learning surgical procedures that were otherwise well performed.

In conclusion, the current work shows that 24h of SD affect the topology of the brain network making it less integrated. The less integrated brain network causes selective damage to the attentional function but not to the switching ability. This is probably related to the fact that the stimuli of the two different tasks are processed in different ways through bottom-up and top-down processing, which are affected differently by sleep deprivation.

## Competing interest

All authors declare no competing interest.

## Notes

### Competing Interest Statement

The authors have declared no competing interest.

### Summary of Updates

the title has been changed. an update has been made to the "statistical analysis", "results" and "discussion" sections. Fig. 7 concerning correlations has changed following the new statistical analyzes carried out

